# TRapping and IMaging (TRIMing) of Cells / Multicellular Organisms in Free Living Environment Enabled by Adaptive Lightsheet Optical Tweezer (aLOT)

**DOI:** 10.1101/2024.05.02.591710

**Authors:** Neptune Baro, Jigmi Basumatary, Neeraj Pant, Partha P. Mondal

## Abstract

To be able to trap and image in a live cell / organism on the go is an incredible feat and paves the way for immobilization-free interrogation. This is a step towards the interrogation of cells / live species in their natural environment. To facilitate, a *TRIMing* technique primarily based on an adaptive lightsheet optical tweezer (aLOT) system is proposed. The TRIMing technique combines the benefits of touch-free optical tweezing and high-resolution imaging. The entire system is built on a single platform for rapid interrogation of freely moving live biological specimens. The trapping system combines an electrical-tunable lens (ETL), cylindrical lens, and an objective lens to generate adaptive PSF. The ETL (in the beam-expander) adaptively changes the beam cross-section (to either a parallel beam or converging point-beam) entering the back-aperture of cylindrical lens, resulting in a point or a line spot at the focus. An objective lens placed at the focus of a cylindrical lens converts the spot to a tightly focused diffraction-limited lightsheet or point PSF. Depending on the object type (spherical or elongated), the system can flip between point and sheet PSF at a rate of 200 *Hz*. The system is integrated to a separate fluorescence arm to enable the imaging of trapped objects (cells or organisms). The *TRIMing* system operates in a brightfield mode to optically trap using point / sheet PSF and subsequently switched to fluorescence mode for imaging. The potential of the system is demonstrated by trapping live specimens (HeLa cells and C. elegans labelled with Bodipy dye) and imaging them in a freely moving environment. Characterization shows a point and sheet PSF size of, 43.42 *μm*^2^ and 70.5 *×* 4.9*μm*^2^ with a trap stiffness of 1.15 *×* 10^*−*3^ *pN/nm* and 0.46 *×* 10^*−*3^ *pN/nm*, respectively. Fluorescently-labelled live specimens were investigated that showed the random distribution of organelles (lipid droplets) both in cells and C. elegans. The *TRIMing* system demonstrated a resolution of *<* 0.7*μm*, a contrast of ≈ 0.84, a SNR of ≈ 11 *dB*. This allows a good combination of rapid trapping and high-quality imaging. In addition, the system allows near real-time determination of critical biophysical parameters, such as organelle size of 1.01 *μm* (in cells) and 1.29 *μm* (in C. elegans) with a density of 0.021#*/μm*^2^ and 0.039#*/μm*^2^, respectively. The number of lipid droplets are found to be nearly double for C. elegans as compared to HeLa cells. These parameters are directly linked to the physiological state of live biological species. Overall, the developed *TRIMing* system allows high-quality imaging of live specimens in a free living environment.

**Statement of Significance:** The ability to image live specimens in a free-living environment is phenomenal. The existing techniques often constrain/fix/anesthetize these organisms to image their physiological state. This comes with a lot of conditioning and directly affects the physiological state or developmental process in biological species, especially the brain undergoing neuronal activity. The proposed *TRIMing* technique elevates this requirement by optically trapping the moving object and simultaneously imaging the internal organelles with high resolution in a free environment. The technique is expected to have widespread applications in diverse disciplines ranging from fundamental cell biology to optical physics.

## I. INTRODUCTION

Manipulating objects using light has fascinating applications, ranging from physical to biological sciences. A system that is capable of both trapping and imaging a live biological specimen is of immense potential. Often this comes at a price of constraining / immobilizing the live specimen for carrying our imaging / investigation. This is not acceptable for studies that require local physiological investigation and related functions including brain imaging. The existing system immobilizes the live specimen (e.g., C. elegans) in a variety of trapping devices or anaesthetizes to image organelle / organ function. This way of imaging live specimen clearly alters / modifies the target function, and makes the investigation more challenging. So, a system that can both trap and image objects in free moving environment is of utmost need for a range of physiological / developmental studies.

Since its first realization in the year 1970 by Arthur Askhin, optical trapping / tweezing techniques has grown into a major investigation tool across sciences and engineering [1]. Subsequent years have seen its application in biology with the trapping of bacteria and red blood cells [2] [3]. This is followed by a plethora of applications in diverse disciplines ranging from fundamental physics to single molecule biology. Some of the unique studies involve, rotation of microscopic objects, short-range colloidal interactions, cellular-liposome interactions, liposome biomechanics, single molecule dynamics [4] [5] [6] [14]. Several variants have evolved over time depending upon the application at hand such as, ring-vortex traps, shape-phase holography, scanning point traps, beam shaping and hyperbolic meta-materials [7] [9] [10] [11] [12] [13]. Off late, a new kind of optical tweezer (LOT) is developed which is primarily based on light sheet illumination rather than the existing point-illumination [15]. All these techniques have advanced the field of optical trap and greatly aided to the measurement of feeble forces in multiple applications [16] [17] [18] [19].

Techniques that facilitate trapping and imaging of living specimens in free moving environment are of great relevance to biology and physics. Often live organism imaging is achieved by immobilizing it to surface by glue or constraining its motion in space or by anaesthetizing it. Existing techniques used for immobilizing live specimens use agar-pads, where worms are kept on a thin slab of agar and by placing a coverslip on it [20] [21]. Other prevalent techniques use anesthetics (sodium azide or levamisole) to affect immobilization [23] [22]. Although these techniques are useful, they are known to alter the target function. For example, agar-pad immobilization is invasive and known to disrupt or arrest development, thereby interferes with the biological process under investigation [24] [25]. In this regard, a recent development by Berger et al have demonstrated a novel microfluidic imaging method that allows parallel live-imaging across multiple larval stages using an array of microfluidic trap channels [26]. Moreover, confining larval movement to microchambers with fast imaging has shown the potential for *>* 48 *hrs* of developmental study (from larva to adult stages) [27]. In another application, a hydraulic valve is used to immobilize live organisms (such as, C. elegans) during imaging and left free to move at other times [28]. This technique enables long time imaging. In the same line, long-term mechanical immobilization of organism is achieved using an array of tapered channels, facilitating glue-free imaging. Most of these technique are complex, necessitates expertise and alters the physiological state of the organism. In this regard, a touch-free immobilization / trapping technique along with high resolution imaging may go a long way in studying these organisms.

In this article, we present a new technique termed as, *TRIMing* where a live organism / cell is trapped by an optical force (in a point or lightsheet trap), and interrogated / imaged, and then released, thereby allowing imaging of cells / organisms in its natural environment. Depending on the shape of object, appropriate PSF (point or lightsheet) can be chosen. Since the trapping is achieved without any mechanical means and is essentially touch-free, the technique is rapid for interrogation / imaging. Such a system overcomes the constraints related to constriction in a narrow channel or gluing or anaesthetising as the current practice is. The developed *TRIMing* system is expected to advance cell and developmental biology to the next level.

## II. RESULTS

### A. Lightsheet Optical *TRIMing* System

The schematic diagram of the optical *TRIMing* system is shown in Fig.1. The entire system broadly consists of four main sub-systems: optical trap, fluorescence microscopy, transmission imaging setup and optical detection. The trap sub-system use a 1064 *nm* laser, adaptive beam-expander (including ETL), cylindrical lens (CL), and a high NA (1.25) objective lens to realize a stable optical trap. The electrical tunable lens (ETL) is a special type of lens that can change its aperture to effect convergence / divergence of the output beam, allowing a change in the lens’s focus. The beam-expander expands the beam and directs it to the cylindrical lens. Note that, the case where beam fills the back-aperture of CL (parallel beam) gives rise to a line whereas, under-filled beam (converging beam) produces a point. Since the cylindrical lens focuses light in one direction, the line beam profile at the back-aperture of a 100X objective lens (NA 1.25) produces a diffraction-limited PSF at its focus. The resultant PSF can be switched between point and lightsheet using an appropriate ETL driving current (see, inset in Fig. 1). The second sub-system is essentially a high resolution inverted fluorescence microscope that shares the same objective that is used for trapping and the light source is a 488 *nm* laser. Transmission white light illumination is the third sub-system that uses white light source with a 10X objective lens for transmission imaging. Finally, the detection sub-system consists of a sensitive *sCMOS* detector placed in the 4*f* detection setup for capturing high-resolution images (for both, transmission and fluorescence module). Again, the same trapping objective lens is used for collecting the emission for the 4f detection sub-system. Specifically, the images are recorded by a sensitive camera with a quantum efficiency of *∼* 84% (sCMOS, Andor Inc., UK). A combination of notch and bandpass filters are used to filter out background and illumination light.

**FIG. 1:**
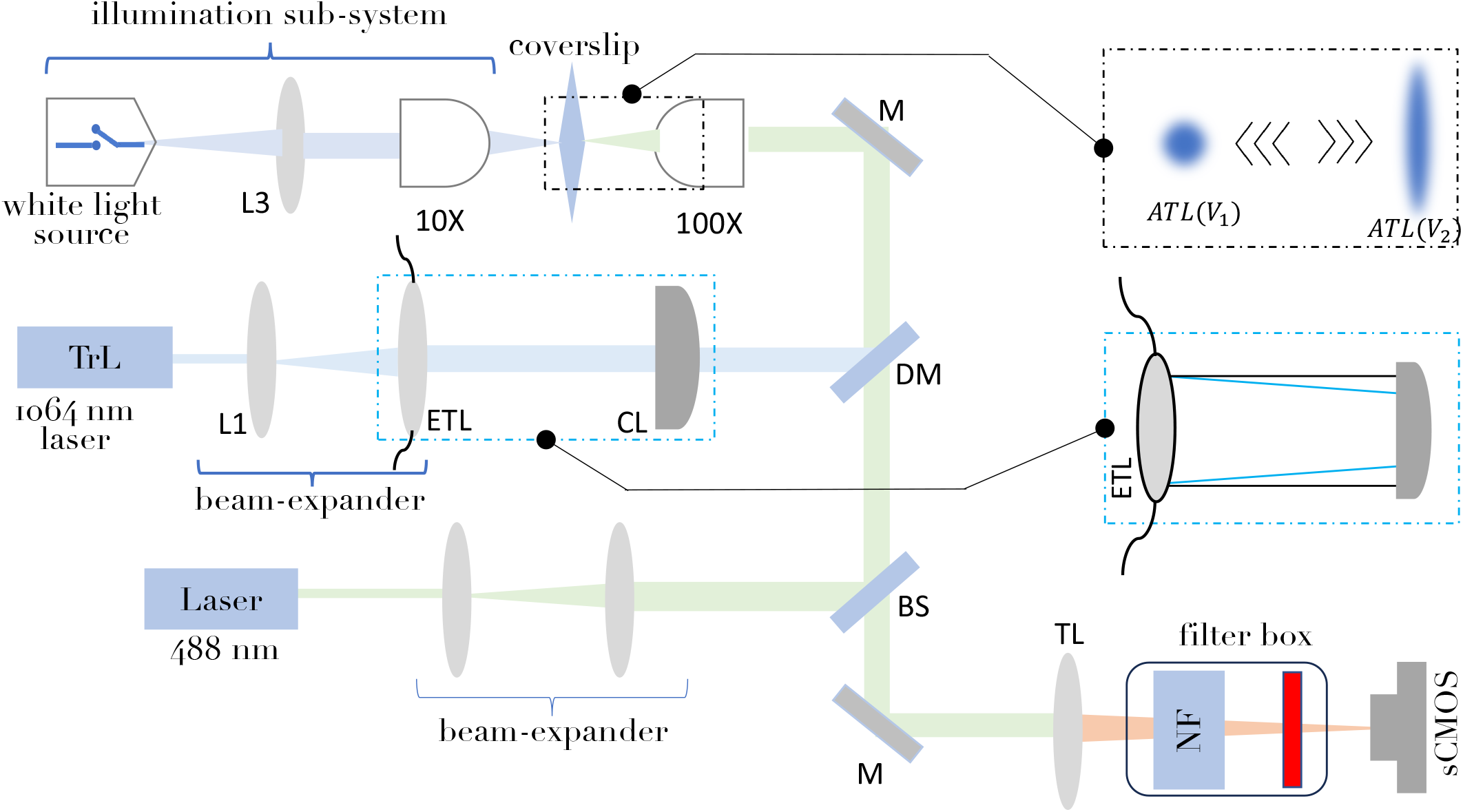
Schematic diagram of aLOT system : The system consists of 4 major sub-systems. The trapping comprises of 1064 trap laser (TrL), ETL, cylindrical lens (CL), 805 *nm* short-pass dichroic mirror (DM) and high NA 100X objective lens. Fluorescence imaging consists of a 488 nm excitation laser, beam-expander and 100X objective lens. A transmission imaging system that consists of white light source, and 10*X* objective lens. Finally, the detection system that comprises 100X objective lens, multiple mirrors, tube lens (TL), filter-box (1064 *nm* notch filter NF, 750 *nm* low-pass filter and 520 *±* 50 band-pass filter) and a sensitive *sCM OS* camera.

### B. Characterization and Calibration of *TRIMing* System

To access the benefits of *TRIMing* system, it needs to be calibrated and the PSFs require characterization over the variability of focal-lengths offered by ETL in the trapping sub-system. To characterize the trap PSF, the *TRIMing* system is operated in the reflection mode as shown in Fig. 2A. Depending on the current supplied to the ETL, the resultant PSF can be switched between point-PSF and sheet-PSF. Fig. 2(B1,B2) shows the characteristics of system PSFs with ETL current and alternately with ETL focal-lengths. Since the curvature of ETL lens aperture decreases at low currents (*<* 88 *mA*), this resulting in longer focal length and near-parallel beam at short distances from the cylindrical lens aperture, nearly filling the back-aperture which after focused by the cylindrical lens produces a lightsheet PSF. However, the curvature increases at large current values (*>* 190 *mA*), resulting in short focal length (or converging beam) producing a point focus passing through the central region of the cylindrical lens. This pencil of parallel beam travel through the central region of cylindrical lens as it is (without deflection), which upon further focused by high NA objective lens produces point PSF. Note that the pencil of beam is broad enough to fill the back aperture of objective lens. So, depending on the ETL current, both point and lightsheet PSF can be realized and toggled at a rate of 200 *Hz*. The measurements of the PSFs are carried out in fluorescence mode using fluorescent gel-block as a sample (see, Methods section). The corresponding PSFs are shown in Fig. 2(B3,B4). The dimension of point PSF is calculated to be (*X, Y*) = 6.68*μm ×* 6.5*μm*, whereas the size of light sheet PSF is, (*X, Y*) = 70.5*μm ×* 4.9*μm*. This adds versatility to the *TRIMing* system by allowing trapping of both spherical as well as elongated objects.

**FIG. 2:**
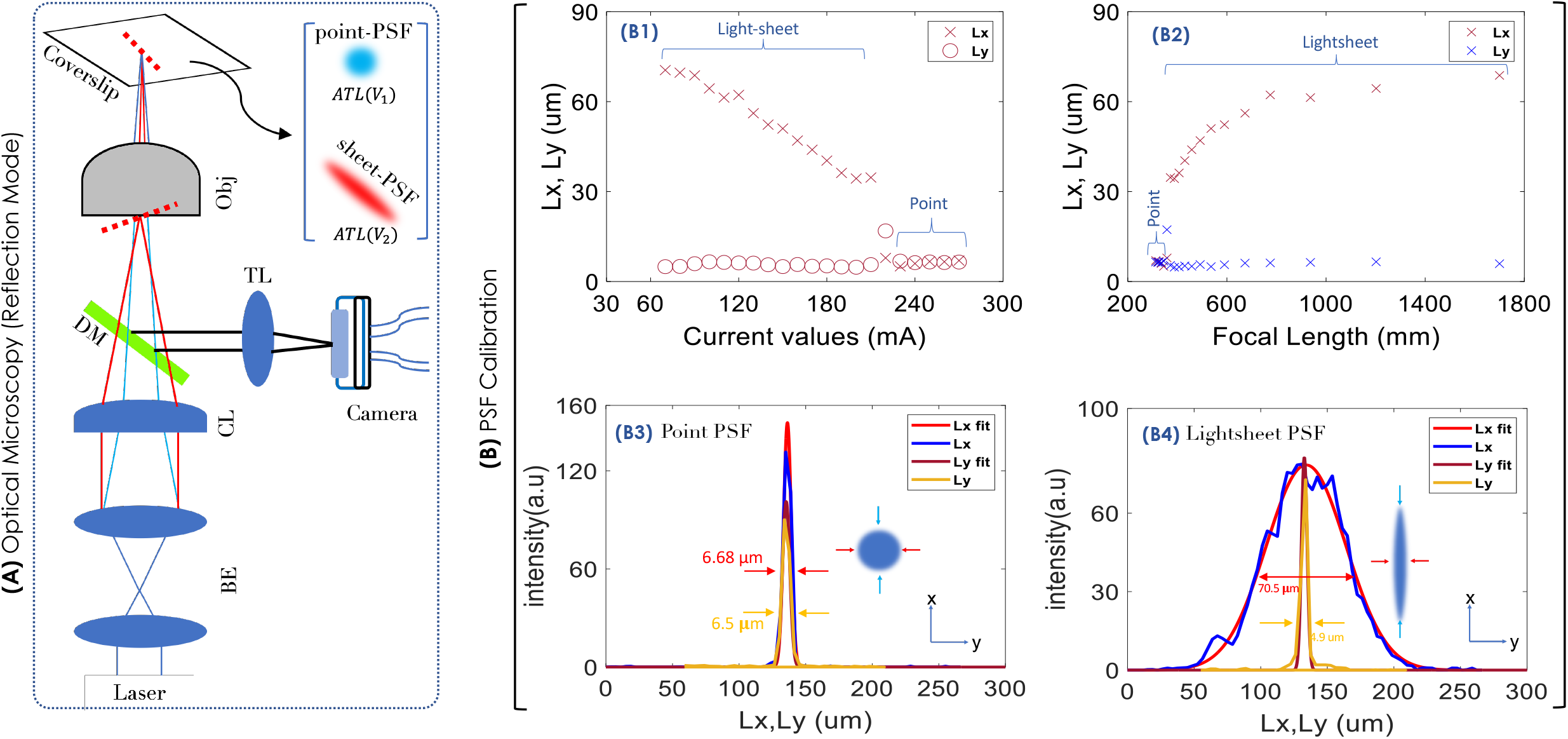
Characterization of system PSFs: (A) Schematic diagram of the optical setup used for characterizing the system PSF. (B1, B2) PSF dimension along X and Y axis as it toggles between point-PSF and sheet-PSF at varying current / focal length values. (B3, B4) The size of experimental point-PSF, and lightsheet-PSF along X and Y axes.

### C. Trap Stiffness and Force Calculations

Trapping an object directly depends on the strength of an optical trap and the force it imparts on the object. Indirectly, this implies that transparent objects with less mass are strongly attracted towards the trap-center and vice-versa. To determine the stiffness and other calibrations (related to stage calibrations), dielectric silica beads (size *∼* 2*μm*) suspended in deionized water was used. A drop of the bead solution is dropped on the coverslip which is placed on the oil immersion objective lens (Olympus, 1.25 NA, 100X). The beam is focused in the solution and beads are trapped at the working distance of the objective lens (along *z*-axis). The sample stage is further illuminated by white light based transmission imaging sub-system, and the beads are visualized using a dedicated 4*f* detection system. In the transmission imaging mode, the filters used in the detection allow white light (broad wavelength band) to pass through and reach the detector (CMOS camera). Observation shows that free beads exhibit Brownian motion. When these beads comes in close vicinity of the attractive trap-potential, it dragged towards the trap-center / beam focus. The high speed CMOS camera (operated at 70 fps) captures the bead images from the beginning / free state (exhibiting Brownian motion) to the end / trapped state (i.e, at the trap-center). This allows calculation of travel-time (t) from the number of frames between the initial to final position. Based on this, the trap-stiffness is given by, *k* = 6*πηr*_*b*_*/t* [15]. Here, *η* is the viscosity of the suspension medium, *x* is the displacement from beginning (free state) to the end (trap-center). *r*_*b*_ is the bead radius, and *t* is the travel-time. Fig. 3A shows the variation of trap-stiffness (*k*) with ETL current value. Depending on the ETL current, either of the traps (point-trap or lightsheet trap) is realized. At low currents, ETL focus is large, producing near parallel beams at short distances from ETL, which upon focused by cylindrical lens produces lightsheet. On the other hand, high current results is short focus and the resulting converging beam passing through the central region of the cylindrical lens goes through as it is without causing any lensing effect, giving rise to a point-focus. This results in point-PSF at the working distance of high NA objective lens. For our case, lightsheet-PSF is observed for *I* ∼ [80 *−* 150 *mA*], whereas, point-PSF is observed at currents, *I* ∼ [250 *−* 290 *mA*]. The corresponding force versus current characteristics is shown in Fig. 3B. The forces realized by the trap range from, 0.39 *pN* for lightsheet trap to 1.15 *pN* for point-trap. So, lightsheet traps are 3 times weaker than point-traps. This is due to the size of lightsheet as compared to point over which the photons are distributed. This overall suggests that the lightsheet-trap has a large region of influence as compared to point-trap but proportionately less stiffness and force. However, this can be compensated by increasing the laser power.

**FIG. 3:**
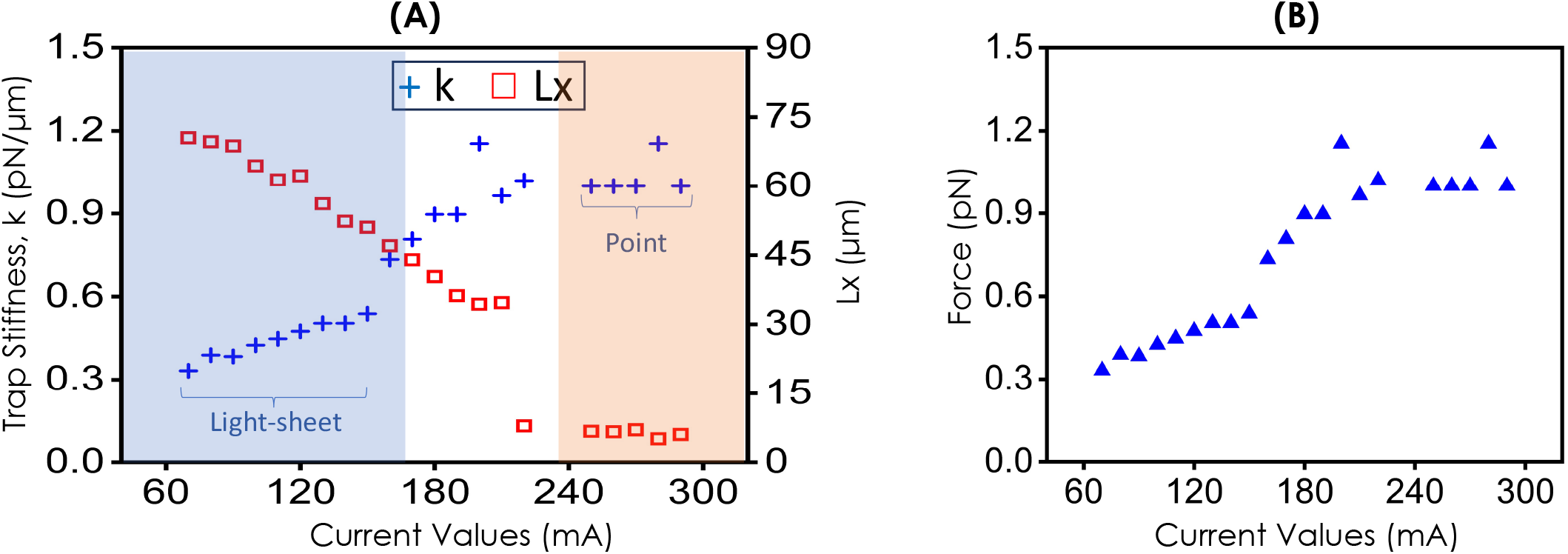
Optical Trap Stiffness: (A) The adaptive evaluation of trap-stiffness with ETL current for point and lightsheet PSF along x-axis, showing ∼ 3 times weaker stiffness for lightsheet trap as compared to point trap. (B) The corresponding force on the particle with ETL current.

### D Trapping HeLa Cells and Multicellular Organisms (C. Elegans)

To validate the working of *TRIMing* system, objects of various shape and size are trapped. We choose to trap microscopic dielectric silica beads (spherical, size *∼* 2*μm*), live HeLa cells (nearly spherical, size *∼* 26*μm*), and elongated live multicellular organism (C. elegans, about 240*μm* long with a diameter of 9.5*μm*). Trapping objects of various shape and size is enabled by adaptive PSF (point / lightsheet). To demonstrate, live HeLa cells and C elegans are trapped while they are in a free environment. The cells were suspended in cell media (PBS), while C elegans are kept in a culture medium (M9 buffer). The point-trap is generated at an ETL current of 259 *mA*, while elongated lightsheet-trap is realized at a current value of 70 *mA*. Fig. 4 shows the respective PSFs and its location. HeLa cells (in PBS) and C.elegans (in M9 buffer) are suspended in media and thoroughly pipetted. The solution is dropped on a coverslip, which is attached to a 3-axis nanopositioning stage. Initially, a point focus is generated near the coverslip surface for trapping dielectric beads and HeLa cells. Fig. 4A shows time point of dielectric beads in the process of being trapped. The corresponding images of the trapping process was captured using a high-speed *CMOS* camera operated at 70 fps. The time taken for the entire journey (from free position to trap-center) took a time of 0.55 *secs*. Subsequently, HeLa cells are trapped by point trap as shown in Fig. 4B and the images are captured at regular intervals during the process. The entire trapping process is captured in **Supplementary Video 1**. On an average, HeLa cells took 2.22 *secs* to reach the trap-center. Subsequently, the point-PSF was switched to lightsheet PSF for trapping C. elegans. Fig. 4C shows acquired images during the process of C. elegans being trapped by lightsheet-trap. The corresponding process is shown in **Supplementary Video 2**. C. elegans took a time of about 10.43 *secs* to reach the trap-center. The relatively longer time is due to the mass of C. elegans as compared to dielectric beads and HeLa cells. A graph showing the comparative time taken by the objects (dielectric beads, Hela cell and C. elegans) is shown in Fig. 4D. Overall, this suggests the efficient trapping of live specimens by the optical trap generated by *TRIMing* system.

**FIG. 4:**
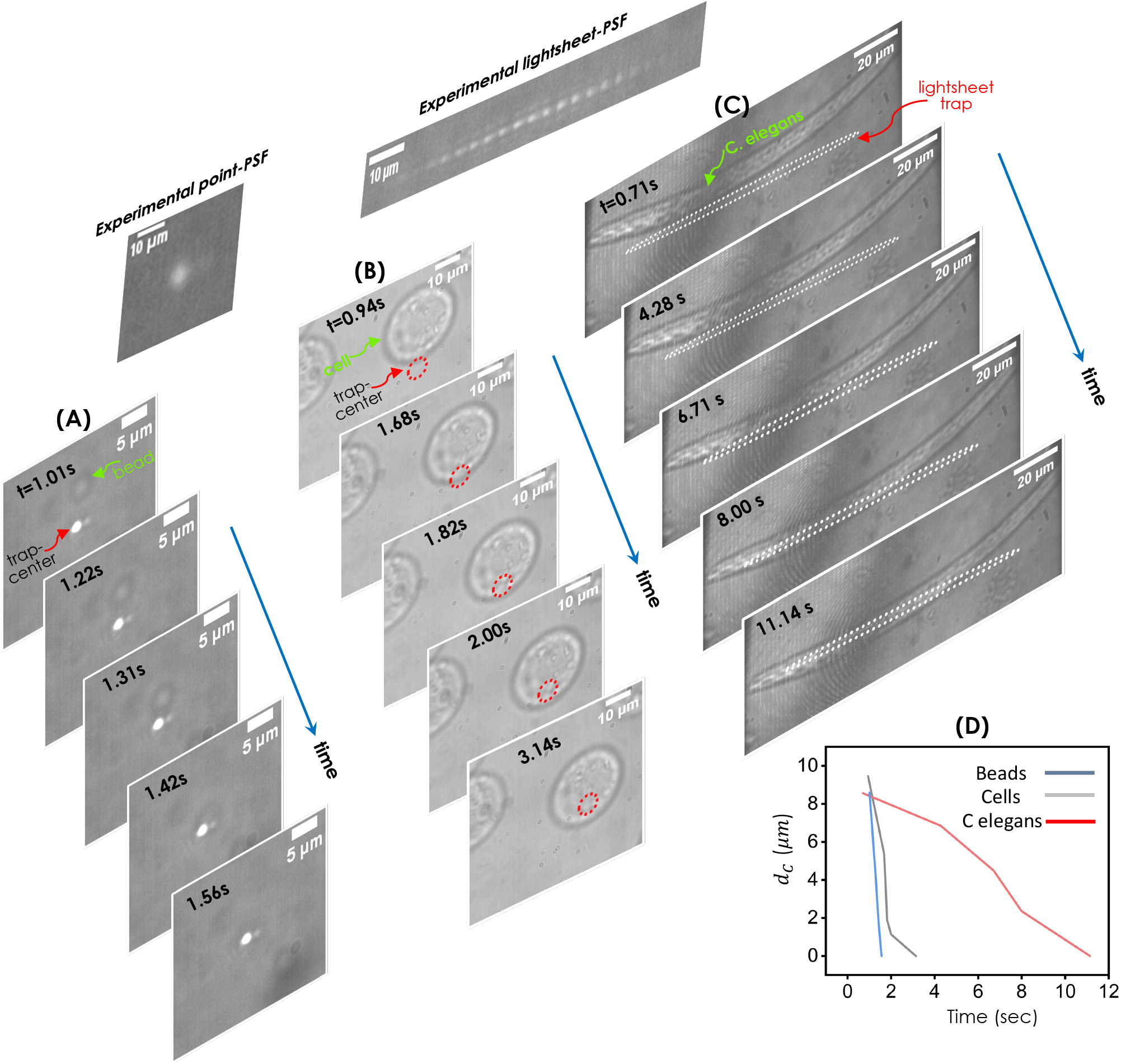
Trapping dielectric beads, HeLa cells, and C. elegans. Recorded point-PSF and lightsheet PSF in the reflection mode. (A) Dielectric bead (size, 2*μm*) being attracted by the accelerating potential. (B) A HeLa cell in the process of being attracted by the potential. (C) Multicellular organism C elegans in the process of being attracted by the potential. The trap-center and bead is indicated by red and green arrow. The corresponding time points are also mentioned for each case. (D) The distance (from free motion to trap-center) versus time taken plot suggesting that large massive objects took more time. The process is shown in recorded **Supplementary Videos 1 and 2**.

### E. *TRIMing* (Trapping and Imaging) On The Go

*TRIMing* necessitates both trapping and imaging of objects on a single platform. This helps rapidly scrutinize and interrogate a large population of species (cells or multicellular organisms) on the go. At the system level, this translates into the integration of a dedicated fluorescence imaging arm (see, Fig. 1). The process begins with the realization of desired PSF for trapping the target object. Depending on the shape of object to be trapped, the PSF (point or lightsheet) is generated. The intended live specimen is first trapped and then imaged.

Fig. 5 shows the time-lapse images of HeLa cells and C. elegans during the trapping process. The entire trapping process can be visualized in **Supplementary Videos 3 and 4**. The images are captured in fluorescence mode with 488 *nm* light for excitation. Both HeLa cells and C. elegans were labelled with Bodipy (*λ*_*ex*_ : *λ*_*em*_ | 493 *nm* : 503 *nm*), which tags lipid droplets. The corresponding protocol for labelling the specimens is discussed in Methods section (see, section III.B). Subsequent to trapping, the object is exposed to fluorescence excitation and the images are recorded by *sCMOS* camera. Since the trapped object (cell or C. elegans) moves from off-focal planes to the focal plane (trap plane), the image resolution (measured in terms of FWHM) changes from 1.245 *μm* to 0.876 *μm*. The same is true for C. elegans for which the image resolution is from 1.530 *μm* to 1.238*μm*. Post imaging, the object is released and the process in repeated for other live objects in the free moving environment.

**FIG. 5:**
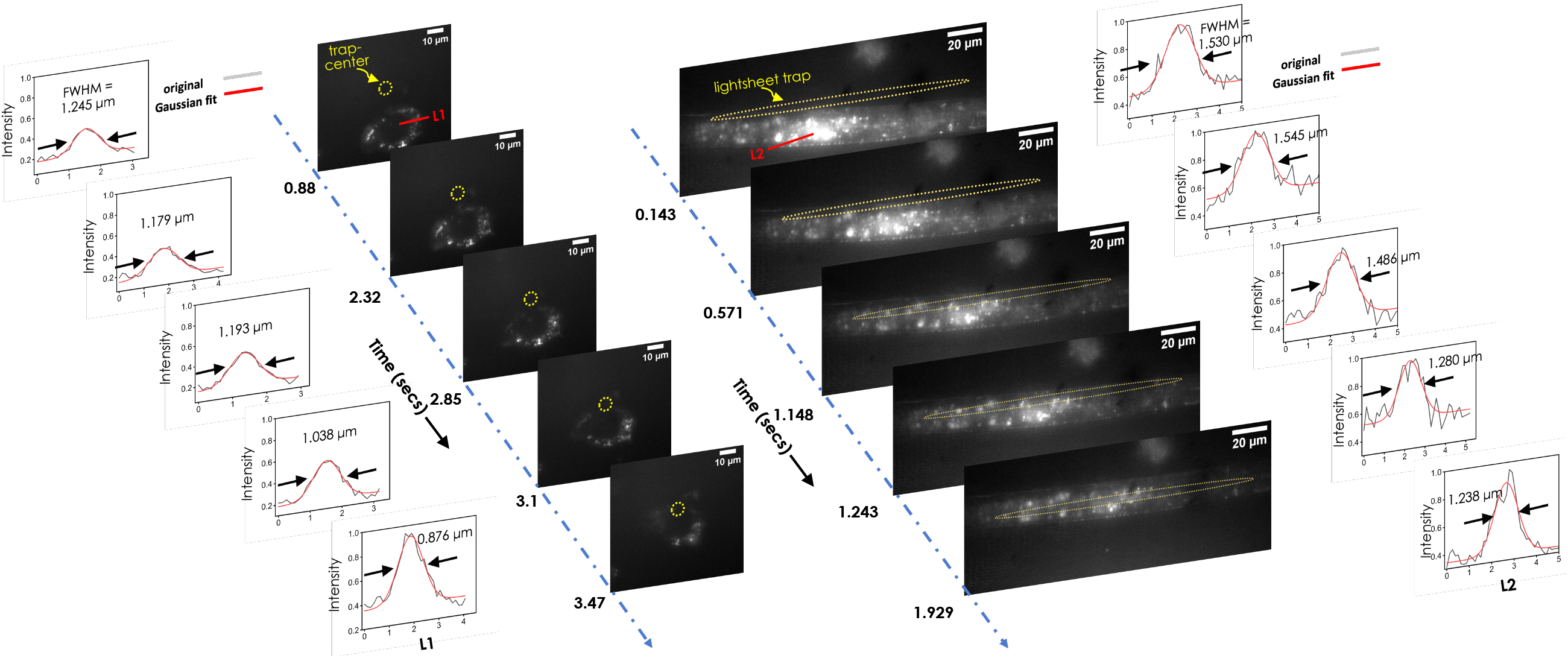
*TRIMing* : (A) Cells are being trapped and imaged during their journey to the trap-center. Alongside intensity plots along red-line *L*1 is also shown that suggests slightly better resolution at trap-centre. (B) Images of multicellular organism (C elegans) in the process of being trapped and imaged. Alongside the time taken to reach the trap-center is also mentioned. The entire process is shown **Supplementary Video 3 and 4**.

### F. Biophysical Parameter Estimation

One of the key purpose of *TRIMing* is to determine critical biophysical parameters related to the object under investigation. Towards this goal, the parameters were estimated on the go in real-time from the recorded images. Fig. 6A shows fluorescence image of C elegans and HeLa cells captured during *TRIMing* process. The corresponding imaging parameters derived from the captured images are also determined. Plots show impressive image resolution, contrast and signal-to-noise ratio. The details are tabulated in Table. 1. The imaging parameters are quite impressive considering the fact that *TRIMing* process is carried out on freely-moving live specimens.

**TABLE I:** TRIMing system parameters & Estimated biophysical parameters for the live specimen.

| Object Statistics and Imaging Parameters |  |  |  |  |  |
| --- | --- | --- | --- | --- | --- |
| Object | Object Shape | Object Size ($\mu m$) | Image Resolution ($\mu m$) | Image Contrast | Image SNR (dB) |
| Silica beads | point | 2.17 | - | $0.452 \pm 0.09$ | $8.167 \pm 1.01$ |
| HeLa cells | spherical | 26.65 | $\sim 0.89$ | $0.94 \pm 0.01$ | $13.73 \pm 1.14$ |
| C. elegans | ellipsoidal | 240.93 | $\sim 1.44$ | $0.83 \pm 0.032$ | $11.03 \pm 0.28$ |

| Biophysical Parameters (Average) |  |  |  |  |
| --- | --- | --- | --- | --- |
| Object | organelle count | organelle size $A_s$ ($\mu m$) | organelle size statistics $\% > A_s \mid \% < A_s$ | organelle density ($\#/\mu m^2$) |
| HeLa cells | $22 \pm 4$ | $1.01 \pm 0.223$ | 52.94\|47.05 | $0.021 \pm 0.003$ |
| C. elegans | $40 \pm 7$ | $1.29 \pm 0.355$ | 58.06\|41.93 | $0.039 \pm 0.008$ |

**FIG. 6:**
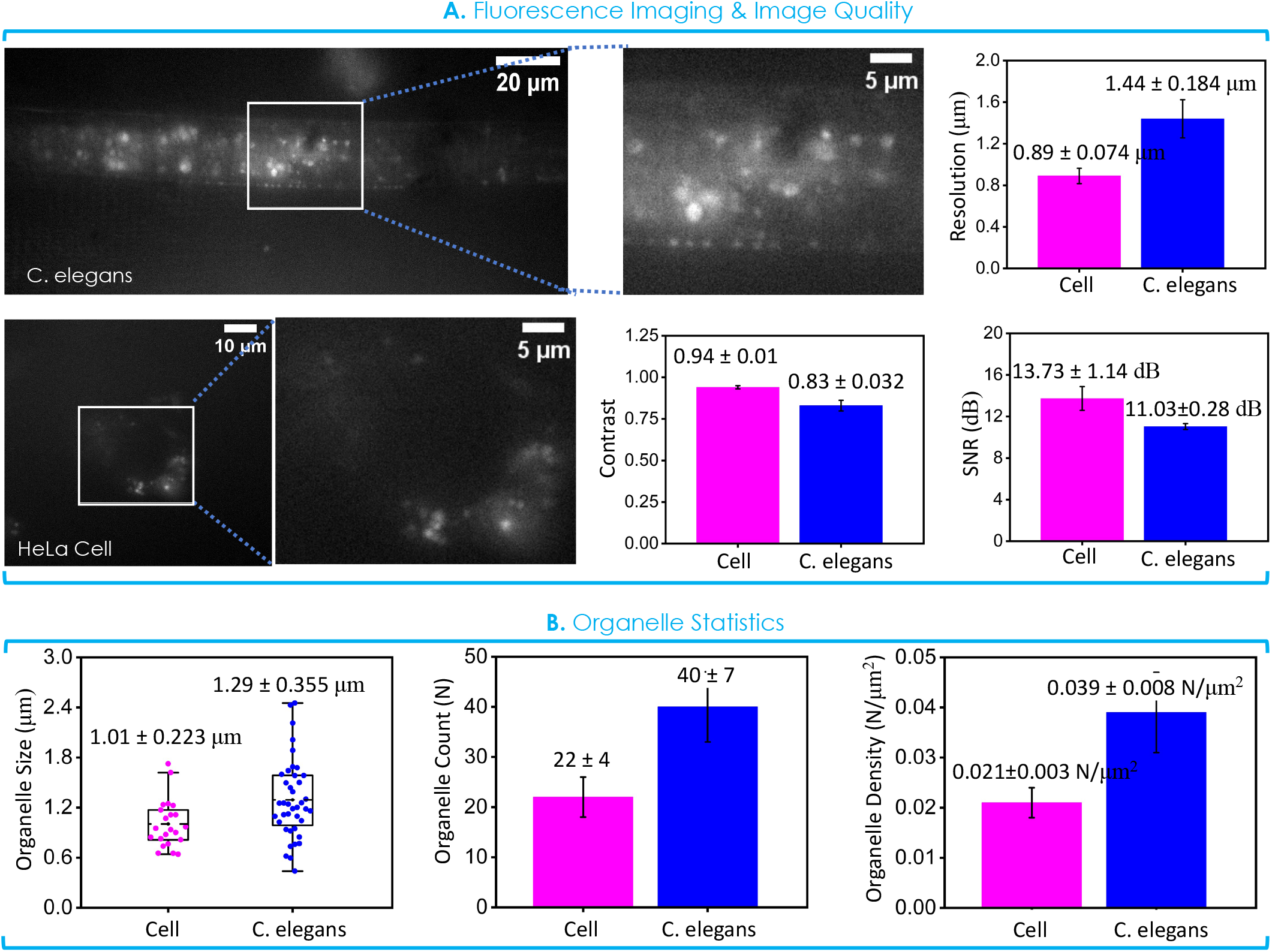
Imaging & Organelle-level Statistics : (A) Fluorescence imaging of multicellular organism (C. elegans) and HeLa cells during the trapped state. The corresponding image quality parameters (resolution, contrast and SNR) are determined on the go. (B) Estimated critical parameters (organelle / organ size, its count and density) of sub-cellular organelles (for cells) / organ (for C. elegans).

Subsequently, critical biophysical parameters related to cells and organisms are determined. Specifically, the number of organelles, its size and density are calculated. For live HeLa cells, the size of Lipid Droplets is found to be 1.01*μm* with a count and density of 22 and 0.021#*N/μm*^2^, respectively. However, for multicellular organism (C. elegans), the parameters are found to be 1.29*μm* (size), 40 (count), and 0.039 #*N/μm*^2^ (density). Overall, TRIMing provides an amazing capability for determining details at the level of sub-cellular organelle / organ of live specimens in free environment. In general, the technique (*TRIMing*) is expected to advance both physical and biological sciences, and developmental biophysics in particular.

## III. CONCLUSION & DISCUSSION

Imaging and interrogating a live cell / organism in its natural environment is an incredible development. Such a possibility will have implications in a range of fields ranging from natural to engineering sciences. In addition, the possibility of contact-less trapping, and manipulation may enable interrogation without the effect of constriction, giving a near touch-free experience. In this work, we report development of a new optical system where both the trapping and interrogation are performed by light, thereby opening a new kind of interrogation system. The technique is performed to optically trap live specimens (cells / multicellular organisms), and the interrogated / imaging is performed non-invasively.

The developed system comprises of four different sub-sections: (1) optical trapping, (2) white light illumination, (3) fluorescence imaging, and (4) 4*f* optical detection. The first subsection use a series of lenses (including ETL) and mirrors to achieve adaptive and fast switchable PSF (either point or lightsheet) (see, Fig. 1). This kind of traps are universal in the sense it can be adapted to trap objects of any shape and size. The other integrated sub-system is essentially a transmission microscope that primarily consists of white light source and a 4f system for visualizing the specimen. The third sub-system is a high-resolution inverted fluorescence microscope for carrying out imaging in tandem with the trapping sub-system. Here the fluorescence system is designed for imaging organelles on the go while the object is in free environment. Two live specimens (HeLa cell and C. elegans) are considered to demonstrate the technology. The entire system is synchronously operated to facilitate trap-visualize-interrogate/image on the go.

The PSF is constructed to facilitate trapping of both symmetric (here, spherical beads or cells) and asymmetric elongated objects (such as, C. elegans). The electrically-tunable lens allow such an adaptive PSF that can rapidly toggle between point to line / sheet PSF. The stiffness of the trap changes rapidly depending on the PSF shape, with point trap generating a stiffness of 1.15*pN/μm* and lightsheet PSF of stiffness *∼* 3 times weaker. Since, the lens focus is controlled electronically, the system PSFs can switch rapidly (at a rate of, 200*Hz*). This gives a better control of trapping a range of objects and rapid interrogation of the specimen of interest.

To validate, two different live species (HeLa cells and C. elegans) are trapped while they are freely moving. This is demonstrated in brightfield mode (where, the trapping is achieved by trap-laser and detection is carried out using white light). The cell images are captured at varying time points, beginning from its position of free state to trap-state at the trap-centre (see, **Supplementary Video 1**). Subsequently, the spherical PSF is switched to elongated PSF to trap C. elegans and the images are collected at different time-points (see, Fig. 4 and **Supplementary Video 2**). This demonstrates the capability of *TRIMing* system to capture particles of different shapes and of different sizes.

In order to achieve realization of complete system i.e, both trap and image on the go, the fluorescence arm is switched on. Again, the specimens (both cell and C. elegans) are trapped and immediately fluorescence image of the specimen is captured (see, Fig. 5, and **Supplementary Videos 3 and 4**). This is incredible considering the fact that both trap and image can be achieved on a single platform without the involvement of any mechanical means. This eradicates the need for constraining (gluing or fixing the sample on a surface) or anaesthetizing the live specimen during interrogation. This is certainly better than the current state-of-the art technique where interrogation / imaging require fixing cells [26] [27]. Moreover, *TRIMing* technique facilitate determination of biophysical parameters related to sub-cellular organelles for HeLa cells / organs for C. elegans (see, Fig. 6). The capability of *TRIMing* as a sophisticated tool is commendable since it allows trapping, imaging and analyzing, all in a single platform.

We anticipate that the availability of organelle-level information such as, number of organelles / organs, its size and density in a freely-moving live specimen will have broad implications in fields ranging optical physics to bioengineering.

## IV. MATERIALS & METHODS

### A. Sample Preparation

#### Silica Beads

Beads based on silicon dioxide in a 5% concentration of 5ml aqueous solution were purchased from Sigma Aldrich, Germany. The bead size is 2 microns with a standard deviation of 0.2 microns. The beads are diluted in deionized water to 1/10th of the original concentration for the trapping experiment.

### C. elegens: Bacterial Food Source and Seeding of NGM Plates

E. coli strain OP50 is used as a food source for the culture of C. elegans. A single colony from an already plated streak is inoculated in an LB media. Inoculated culture is allowed to grow overnight at 37 *°*C. Subsequently, The E. coli OP50 solution is used for seeding NGM plates. The cultured solution can be stored and used for several months. A drop of 10 *μ*L of cultured E. coli OP50 liquid solution is poured on 100 mm NGM plates. The drop is spread using a pipet tip to create a large lawn on the NGM surface. Care is taken to keep the lawn at the centre of the NGM plates. Subsequently, NGM plates were incubated for 12 hours at 37 *°*C to achieve a lawn of E. coli for C. elegans.

### B. Labelling Protocol

#### HeLa Cells

For staining of HeLa cells, we chose BODIPY 493/503 (Invitrogen-D3922), a lipophilic fluorescence that stains for non-polar lipids. Dye stock was diluted in PBS or DMSO to obtain a stock concentration of 2 mg/mL. The stock PBS solution is diluted in PBS to a working concentration of 2 *μ*M. Bodipy binds with the lipid droplets deposited in cell walls.

For staining, Hela cells (Human cervical cancer cell line) were cultured in Dulbecco’s modified Eagle’s medium (DMEM) (Gibco™, Thermo Fisher Scientific, Waltham, MA, USA) supplemented with 10% fetal bovine serum (FBS) and 1% of penicillin-streptomycin. The cells were seeded at a density of 100000 cells in 35 mm dishes and maintained at 37 *°*C in a humidified incubator with a 5% CO2 atmosphere for 24 hrs. The dishes were observed for 70 *−* 80% and washed twice with 1X PBS to remove debris. The debris-free dishes were filled (treated) with Bodipy solution for staining the cells and incubated for 45 minutes at 37 *°*C. A thorough wash with 1X PBS was performed to achieve better SBR (signal-to-background ratio) during an experiment. The cells were detached from the dishes using a diluted trypsin solution (1% trypsin in PBS) and collected in a 1.5 ml Eppendorf tube, followed by centrifugation for 2 minutes at 3000 RPM. The pelletized cells were re-suspended in 1 ml of 1X PBS and used for trapping experiments.

#### C. Elegens

For staining of C. elegans, we use 5 *μM* Bodipy 493/503 (Invitrogen-D3922) solution, which binds with lipid droplets in C. elegans. C. elegans strain N2 was chosen and cultured on Escherichia coli OP50 lawns on NGM agar plates at 25 *°*C for 2 days. Freshly harvested L1 stage C. elegans were taken in a 1.5 ml Eppendorf tube and incubated with bodipy dye (5 *μ*M) for 5-6 h. After staining, it was washed twice with M9 buffer followed by cetriguation at 3000 RPM for 5 minutes. The pelletized samples were re-suspended in 1 ml of M9 buffer and used for trapping experiments.

### C. Image Acquisition and Data processing

For acquisition, a CMOS (Gazelle, Pointgray, USA) camera is employed with a 125 mm tube lens to acquire the brightfield images. The graphical programming environment, Labview (NI instruments), is interfaced with the CMOS camera to control the data acquisition. The camera is adjusted to a suitable exposure time for crispy images and is set to a 1355 x 500 window size for faster acquisition. For trapping of HeLa cells, the ETL is set to the current value of 260 *mA*, and aLOT system is calibrated to achieve the point PSF. Subsequently, 5-minute videos were captured at a frame rate of 70 *fps* to visualize the trapping of Cells. For trapping of C. elegans, the ETL is set to the current value of 70 mA to achieve the LS PSF.

For the acquisition of fluorescence images, an sCMOS camera is employed with a 125 mm tube lens. The images underwent pre-processing and post-processing like Gaussian blur filtering followed by Hough transform for statistical estimation of Biophysical parameters.

### D. Biophysical parameters

The developed system can be employed to trap and image cells and multi-cellular organisms simultaneously. We targeted the lipid droplet cell organelles, which store neutral fats such as triacylglycerol (TAG) and cholesterol ester in C. elegans and cells. We used BODIPY 493/503 (Invitrogen-D3922) to stain lipid droplets in C. elegans and Cells. It binds with the droplets and gives out fluorescence (510 nm) on excitation (488 nm). After a stable trap, the distribution of the droplets in different parts of C. elegans and cells was captured using an sCMOS camera. We chose an image with C. elegans trapped at the LS center and one with cells trapped at the point spot center. The acquired images were subjected to pre-processing, such as Gaussian blur filtering in imageJ. A Hough transform function (available in matlab) is used to determine the size of the lipid droplets in C. elegans and cells. ROIs generated from pre-processed images were fetched to the function, and a statistical estimate was evaluated. It was observed that the average size of the droplets is 1.29 *μ*m for C.elegans and 1.01 *μ*m for Cells. These numbers match the reported sizes of the lipid droplets in Cells and C.elegans. This shows the effectiveness of the system developed, which can be used for trapping and imaging simultaneously.

## Contributions

PPM and NB conceived the idea. NB, JB, NP, PPM carried out the experiments. NB, JB, NP prepared the samples. PPM wrote the paper by taking inputs from all the authors.

## Data Availability

The data that support the findings of this study are available from the corresponding author upon request.

## Supplementary Info

The manuscript is supported by 4 supplementary videos.

## Disclosures

The authors declare no conflicts of interest.

## Notes

### Competing Interest Statement

The authors have declared no competing interest.

